# Flexibility and design: conformational heterogeneity along the evolutionary trajectory of a redesigned ubiquitin

**DOI:** 10.1101/081646

**Authors:** Justin T. Biel, Michael C. Thompson, Christian N. Cunningham, Jacob E. Corn, James S. Fraser

**Author notes:** corresponding author (JSF).

## Abstract

**Summary:** Although protein design has been used to introduce new functions, designed variants generally only function as well as natural proteins after rounds of laboratory evolution. One possibility for this pattern is that designed mutants frequently sample nonfunctional conformations. To test this idea, we exploited advances in multiconformer modeling of room temperature X-ray data collection on redesigned ubiquitin variants selected for increasing binding affinity to the deubiquitinase USP7. Initial core mutations disrupt natural packing and lead to increased flexibility. Additional, experimentally selected mutations quenched conformational heterogeneity through new stabilizing interactions. Stabilizing interactions, such as cation-pi stacking and ordered waters, which are not included in standard protein design energy functions, can create specific interactions that have long range effects on flexibility across the protein. Our results suggest that increasing flexibility may be a useful strategy to escape local minima during initial directed evolution and protein design steps when creating new functions.

## Introduction

One of the major aims of both protein engineering and computational design is to create proteins with new or enhanced functions (Butterfoss and Kuhlman, 2006). Proteins can be redesigned to promote new catalytic reactions or, in the simpler case, for binding to new partners. Recent years have seen successes including the design of new enzymes (Kiss et al., 2013), new small molecule binders (Tinberg et al., 2013), and new protein-protein interfaces (Karanicolas et al., 2011). However, these redesigned proteins are generally optimized with many rounds of laboratory evolution to achieve functions approaching those of natural systems (Blomberg et al., 2013; Khersonsky et al., 2010). There are several ways that the engineering or design process can fall short of optimal function: the protein may not fold in the conformation predicted; the intended structure may not imbue the desired function; or the protein may dominantly populate nonfunctional conformations. We (Bhabha et al., 2015), and others (Korendovych and DeGrado, 2014; Osuna et al., 2015), have speculated that the third explanation is a major reason why redesigned proteins are only marginally functional initially. The hypothesis is that initial mutations introduced during redesign disrupt the native packing of the parental wild-type (WT) protein, resulting in increased flexibility and sampling of nonfunctional conformations.

To improve function, subsequent directed evolution experiments fix mutations that act to stabilize the functionally important conformations found in the broadened ensemble of the redesigned protein (Frushicheva et al., 2014). This pattern of design and selection has been performed for functions including small molecule binding (Tinberg et al., 2013) and protein-protein interaction (Karanicolas et al., 2011), the most developed examples of directed evolution changing dynamics and function emerged from enzyme design. Molecular dynamics simulations of designed enzymes indicate that initial designs suffer from poor preorganization (Frushicheva et al., 2010) and that mutations acquired during further selections reduce conformational dynamics near the active site (Osuna et al., 2015). Molecular dynamics simulations have also been used as a computational filter to screen out designs that suffer from conformational instability (Wijma et al., 2015). In principle, NMR methods could be used to assess whether local dynamics are altered from WT by redesign and subsequent evolution (Johnson and Handel, 1999; Walsh et al., 2001). However, it has remained difficult to track changes in conformational heterogeneity during these processes because of the inability to resolve the relevant conformations in atomic detail for proteins that select for new binding or catalytic activities.

To overcome these limitations, X-ray crystallographic techniques to resolve alternative conformations with atomic resolution have recently been developed (Burnley et al., 2012; Keedy et al., 2015). The key insight enabling these techniques is the recognition that weak signals present in electron density maps represent alternative conformations (Lang et al., 2014, 2010). These signals can be interpreted with a multiconformer model where individual residues are built as a parsimonious set of alternative conformations with differing coordinates, occupancies and B-factors (Woldeyes et al., 2014). These features are often more noticeable in X-ray data collected at room temperature. Interpreting these signals in systems such as CypA (Fraser et al., 2009), Ras (Fraser et al., 2011), and DHFR (van den Bedem et al., 2013), has delivered new insights into the structural basis of correlated protein dynamics and their relationship to function. By monitoring differences in multiconformer models, it should be possible to observe how conformational heterogeneity changes as a function of redesign and laboratory evolution for novel enzymatic or binding activities. Here we investigated the changes to conformational heterogeneity of ubiquitin variants selected to bind tightly and specifically to the deubiquitinase, ubiquitin specific protease 7, USP7, also known as HAUSP in humans (Zhang et al., 2013).

Ubiquitin is a hub protein that binds partners with several different interfaces (Husnjak and Dikic, 2012). Previous studies have linked the conformational flexibility of ubiquitin, particularly in the β1β2 loop, to its ability to bind these diverse binding partners (Lange et al., 2008). Unlike the enzyme design examples discussed above, USP7 is already a natural, albeit weak, binder of WT ubiquitin. Zhang et al (Zhang et al., 2013) subjected ubiquitin to a combination of Rosetta design and phage display to encode additional affinity and specificity for USP7. The premise of the project was to mutate the core of ubiquitin to stabilize the “down” conformation of the β1β2 loop. An initial mutant was selected from a phage display library of 7 sites selected based on Rosetta Design calculations. Originally referred to as u7ub25, this construct is referred to herein as the “core” mutant. The final “affinity matured” mutant, originally called u7ub25_2540, contains an additional 3 surface mutations that were selected by additional phage display experiments (**Figure 1**). While structures of the variants bound to USP7 have not been possible to obtain, we reasoned that the structural and dynamic features of the unbound variants could provide new insights into the forces that stabilize binding. Here, we determined the structures of both the core and affinity matured variants to high resolution using room temperature X-ray diffraction. The structure of the affinity matured variant showed that the β1β2 loop adopted the same “up” conformation as WT ubiquitin (Zhang et al., 2013), and as the minor conformer of the core mutant. Thus, the mechanism by which these mutations lead to increased specificity remains unclear. We found that the heterogeneity of these constructs follow a trend where the initial mutations lead to increased flexibility when compared to previously determined WT ubiquitin crystal structures in different crystal forms and that the final mutations then stabilize a dominant conformation. Our results show how characterizing the conformational landscapes of redesigned proteins could improve protein engineering and computational protein design.

**Figure 1:**
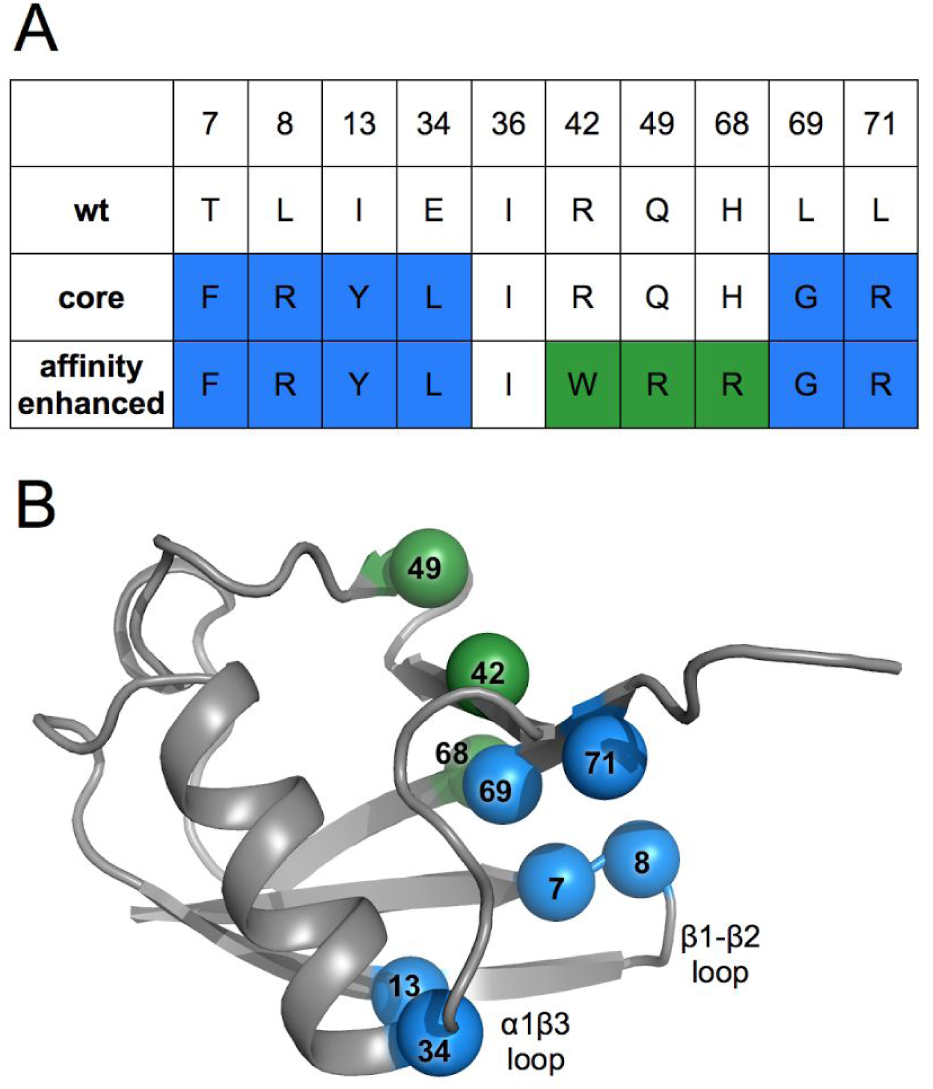
*Locations and identities of mutations made across directed evolution trajectory of ubiquitin*. *A) Table showing amino acid identities of mutation sites for the wild-type, core and affinity matured mutants. Blue coloring represents residue identities first introduced in the core mutant. Green shows new residue identities for the affinity matured mutations.* *B) Wild-type ubiquitin (PDB ID: 1ubq) model shown in grey with spheres representing locations of mutation sites, colored as in panel A and labeled by residue number*.

## Results

### *Conformational heterogeneity of the* β1β2 *loop*

During initial refinement and model building, we noticed electron density clearly motivating the need for multiconformer models (**Figure 2a**). For example, the β1β2 loop displays many large difference density features that cannot all be accounted for by a single conformer model with isotropic B-factors (**Figure 2a**), or even with anisotropic B-factors. In fact, if added first, the anisotropic B-factors can spread into the difference signal for alternative conformations making the density much more difficult to interpret. Therefore, we chose to model alternate conformations first, then add anisotropic B-factors if necessary. In an initial attempt to build a multiconformer model, we used the automated program qFit (Keedy et al., 2015; van den Bedem et al., 2009), which builds multiconformer models via a “sample-and-select” procedure. While qFit has been recently updated to accommodate more backbone flexibility (Keedy et al., 2015), it was not intended to model large displacements with separated backbone density peaks. Consequently, qFit is not able to model highly flexible areas, such as this β1β2 loop (**Figure 2b**). For some residues, such as Phe7 in the core mutant, side chains are even moved by qFit into density that is unlikely to arise from that residue. Clearly the complex backbone shifts in this region are not well captured by these automated techniques.

Due to the limitations of automated model building for this region, we undertook extensive manual interpretation of these regions with alternative conformations. To ensure that the resultant manually built models were consistent with the data, we performed two independent interpretations of the electron density maps (by JTB and MCT, blinded from each other). The approach of comparing independent refinements has been previously used to assess accuracy of structure determination under different purifications (Daopin et al., 1994), with different refinement software (Fields et al., 1994), and with the same data (Terwilliger et al., 2007). Although modeling alternative conformations at low signal levels is necessary to successfully interpret and minimize the local difference density, care must be taken not to interpret signal unless there is a stereochemically reasonable model that can be built (Richardson et al., 2013). The independent refinement procedure allowed us to check for consistent interpretation in regions of high disorder, such as the β1β2 loop, where relevant signals for alternative conformations frequently appear only at low electron density contour levels. The resulting models were almost identical essentially varying only in the interpretation of rotamers for flexible side chains. For residues that had different rotamers or varied in number of conformations modeled (**Supplementary Figure 1**), we evaluated the two models based on rotameric positions, steric clashes, plausible tertiary interactions (e.g. hydrogen bonds to nearby side chain heteroatoms), consistency with the 2Fo-Fc map, and the extent to which local Fo-Fc difference density was explained. After making a consensus model based on these comparisons, we added anisotropic B-factors to protein atoms and finalized the solvent placement. These additions improved the map quality, allowing extra signal for other features to be interpreted in the final model (**Figure 2d**). In the area of the β1β2 loop, the signal at the high and low contour levels clearly defines the molecular envelope, and the difference density in this region is largely reduced (**Figure 2d**). When the final structure is overlaid with the difference density from the single conformer structure (**Figure 2c**), the new additional conformations clearly explain the previous difference peaks.

**Figure 2:**
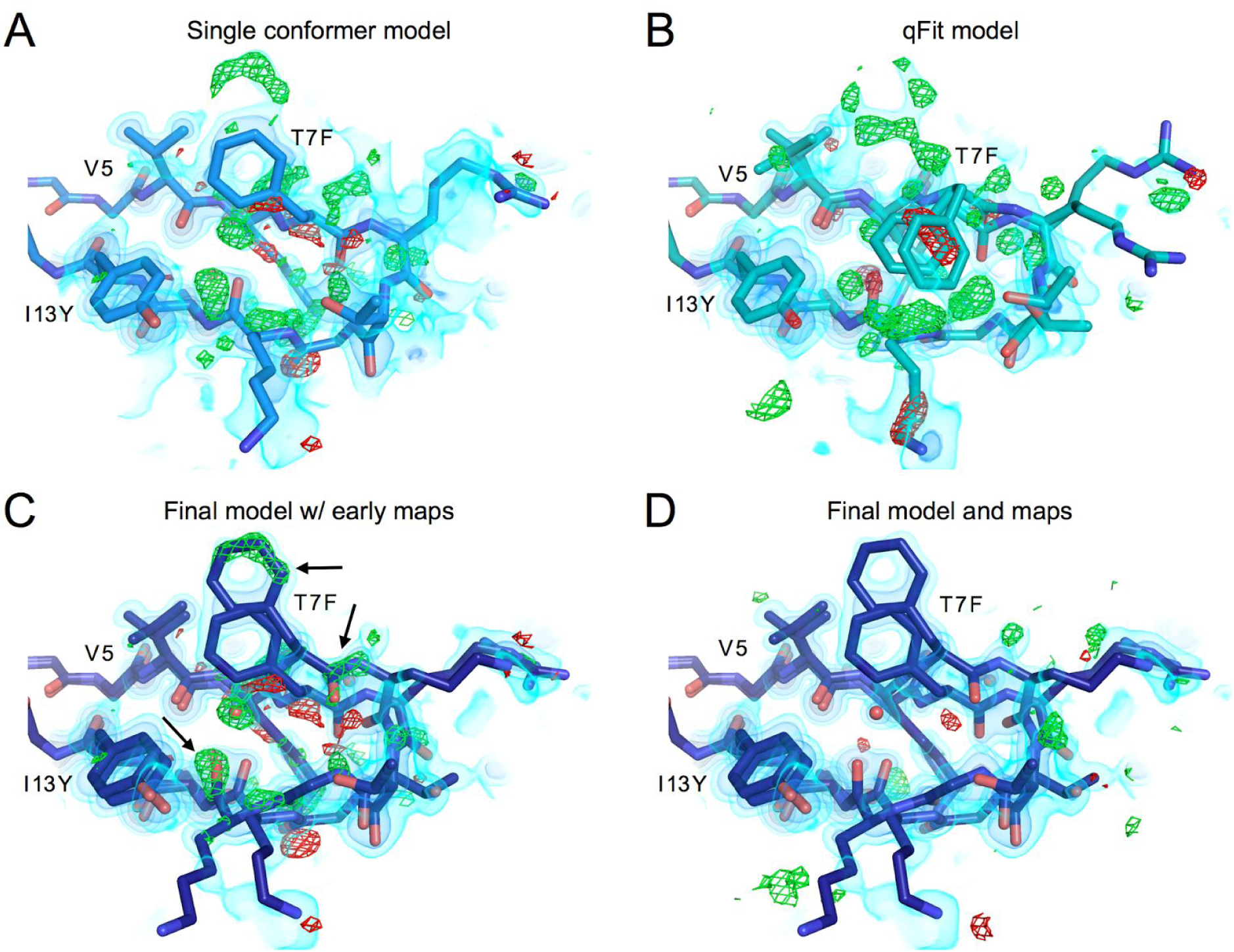
Transcriptome-wide differential gene expression and alternative splicing in ASD. *β1β2 loop of the core mutant showing residues 5-13, models and maps at different points in the refinement procedure.* *A) Best single conformer model and corresponding 2mFo-DFc map shown in volume representation at three contour levels: 0.65 eÅ^−3^, 1.5 eÅ^−3^ and 3.5 eÅ^−3^ from lightest to darkest. mFo-DFc map shown in green and red mesh at 3 or −3 eÅ^−3^ respectively*. *B) Output model from qFit 2.0 built from single conformer model. While qFit is able to accurately model the side chain heterogeneity at the more ordered base of chain for residue 5 valine, qFit was unable to capture the backbone heterogeneity in this loop, and in fact may be mislead by the complex density as can be seen for the clearly misplaced Phe7 side chains.* *C) Final, manually-built multiconformer model with final 2mFo-DFc map, and single conformer difference map (mFo-DFc). This shows how the newly built heterogeneity corresponds to the major difference peaks in the original map*. *D) The final manually-built model with corresponding maps. While some difference features still exist, the difference signal in this region has largely been reduced in comparison to the original single conformer structure*.

The heterogeneity of the β1β2 loop consists of large shifts of the Cα atoms, separating alternative states by as much as 4.5 Å. Alternative conformations of backbone carbonyls are observed pointing in different orientations and the backbone takes two distinct paths for residues 9-11. These loop conformations differ from the expected conformations contained in NMR models of the WT protein (**Figure 3c**), in which the β1β2 loop moves in a hinge like manner, with residues 8-10 moving in unison between “up” and “down” conformations (Lange et al., 2008).

**Figure 3:**
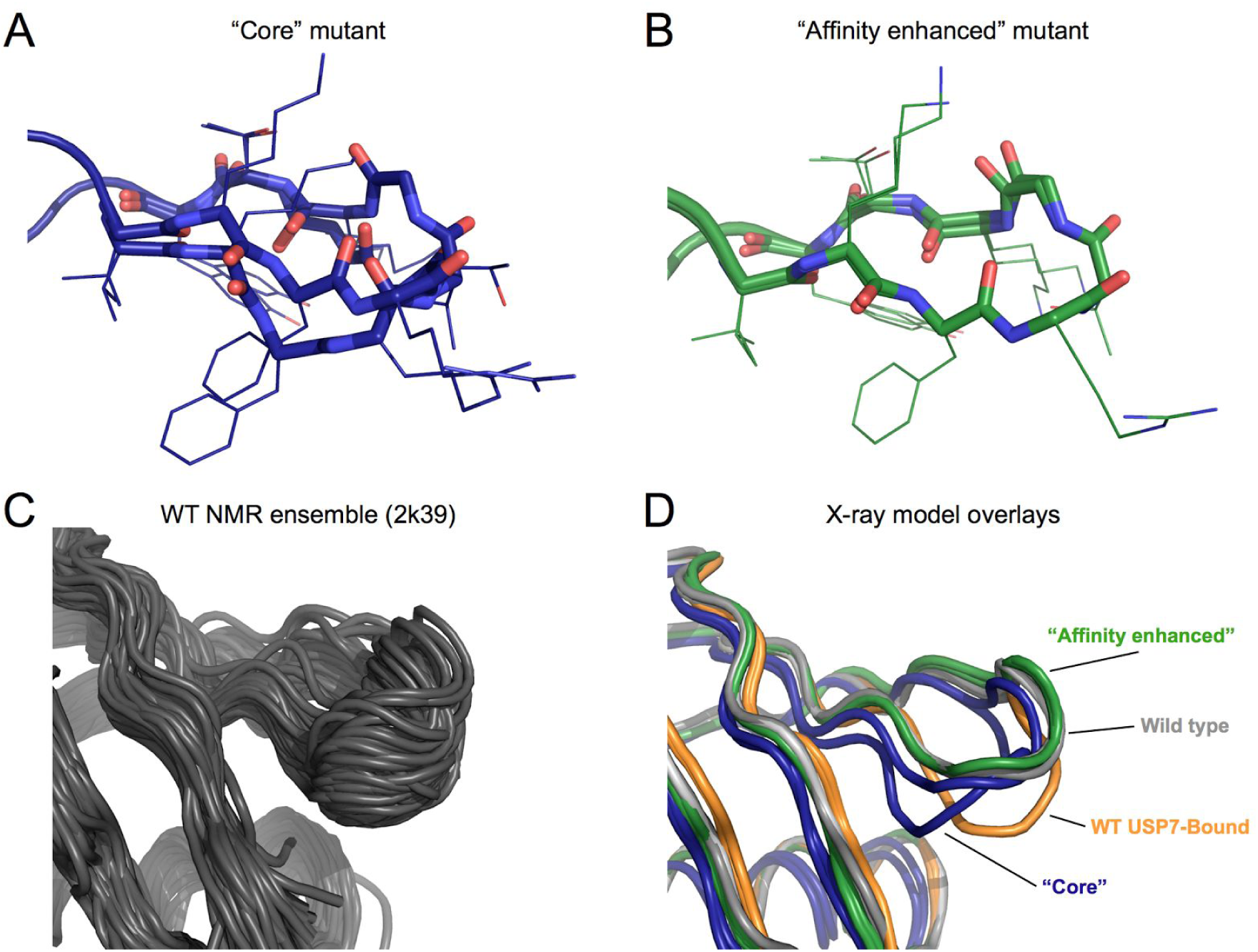
*Conformations of the β1β2 loop* *A) Final multiconformer model of the β1β2 loop for the core mutant construct. Backbone atoms in the loop are rendered in sticks, while the side chains are left as lines*. *B) Final model of β1β2 loop for the affinity matured construct. Panel shown with electron density in Supplementary Figure 2*. *C) Backbone conformational ensemble from NMR relaxation dispersion experiments (PDB ID: 2k39). The majority of conformations exhibit a simple backbone shift, producing a hinge like motion as observed in previous MD simulations*. *D) β1β2 loop conformations from different ubiquitin structures. Wild-type ubiquitin apo-structure (PDB ID: 1ubq) in grey, and bound to USP7 in orange (PDB ID: 1nbf). The core mutant is shown in dark blue, and the affinity matured in green*.

In contrast to the core mutant (**Figure 3a**), the electron density for the affinity matured mutant is consistent with a much less heterogeneous conformational ensemble (**Figure 3b**, **Supplementary Figure 2**). Although the affinity matured conformation is closer to the WT “up” conformation of previous crystal structures than the USP7-bound “down” conformation (**Figure 3d**), it is a more potent and selective binder than either the core mutant or WT (Zhang et al., 2013). Because the crystallization conditions and other considerations such as crystal lattice contacts and data collection temperature are equivalent between these two datasets, our results indicate that the addition of the final three surface mutations in the affinity matured mutant are responsible for quenching this heterogeneity. Below we outline how specific mutations have acted to increase conformational heterogeneity from the WT to the core mutant and to decrease heterogeneity from the core to affinity matured mutants.

### Structural changes of mutations in and near the β1β2 loop

The core variant has 6 mutations that were introduced with the goal of increasing the affinity to USP7, by changing the packing of the protein core to favor the “down” conformation of the β1β2 loop. Two of the mutations are on the β1β2 loop, T7F and L8R, which both mutate the side chains adjacent to the USP7 binding interface. Three of the other mutations are located in the core of the protein (I13Y, E34L and L69G) and the final mutation, L71R, is on the edge of the hydrophobic core on the C-terminus.

Many of the mutations made in the core variant are adjacent to each other, leading to compensatory effects on packing. At the base of β1β2 loop the WT Thr7 is replaced by a bulkier Phe residue, which is then compensated by a large-to-small mutation (L69G) on the adjacent β5 strand (**Figure 4 a,b**). While steric packing is conserved, T7F can no longer make hydrogen bonds to the backbone carbonyl of Lys11 or to the side chain of Thr9. Both residues that previously participated in hydrogen bonds in the WT background show enhanced heterogeneity in the core mutant, as seen for the Lys11-carbonyl (**Figure 4b**). For residue Thr9, the lack of a hydrogen bond allows the threonine side chain to flip out relative to the WT conformation (**Figure 4 a,b**). In the absence of the WT leucine side chain for residue 69 (L69G) for which the newly introduced residue 7 phenylalanine (T7F) to pack against, we observed dramatically shifted conformations of Phe7 in the electron density. Collectively, these features likely stabilize the β1β2 loop in WT ubiquitin, and their absence correlates with the increased heterogeneity of the core mutant.

**Figure 4:**
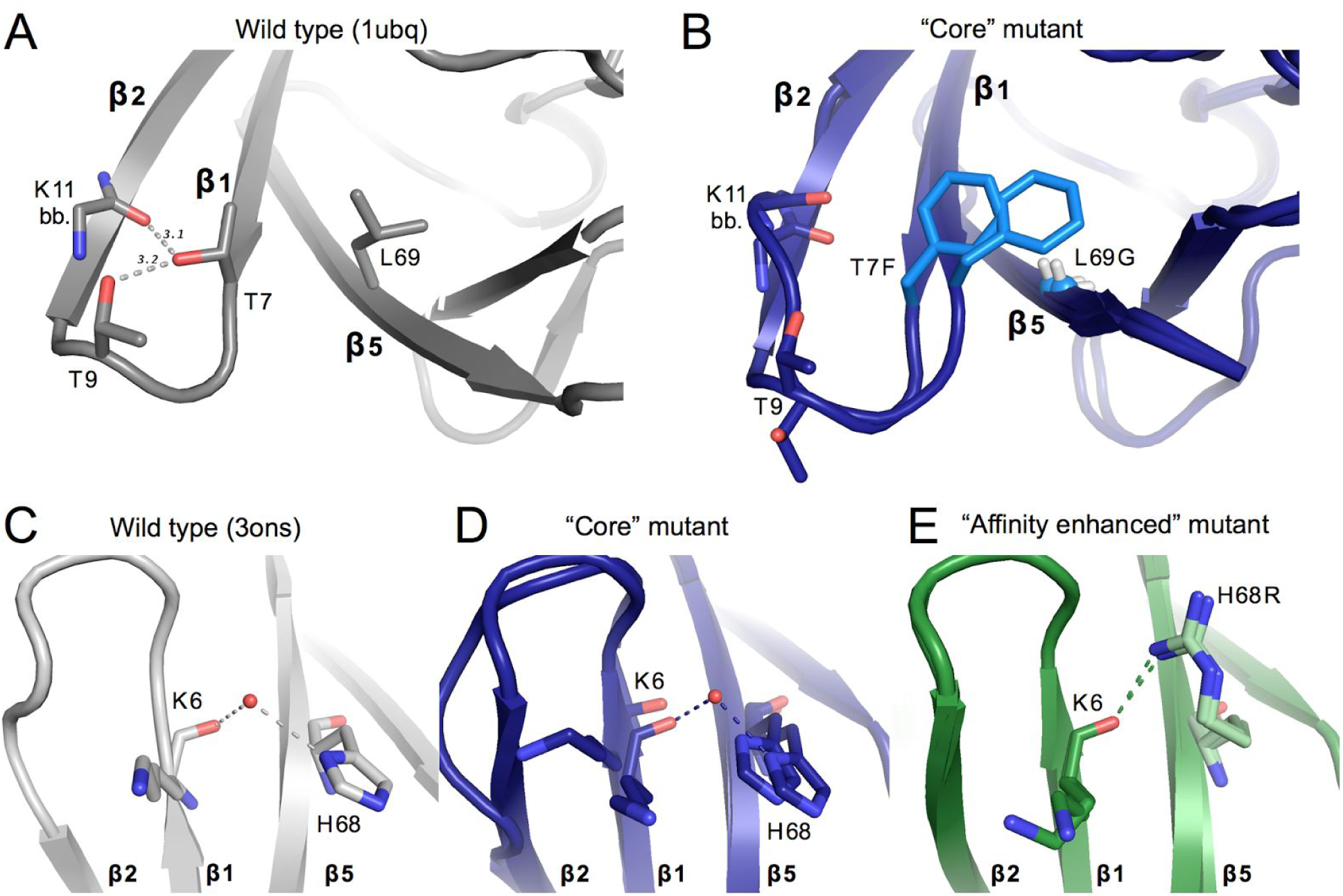
*Structural changes upon mutation near β1β2-loop* *A) Key packing and hydrogen bond interactions around mutation sites T7F and L69G. WT ubiquitin (1ubq) is shown in gray. Gray dashed lines show hydrogen bonds existent in WT ubiquitin between residues 7, 9, and 11. Sticks are shown for the side chains of residues 7 and 9, as well as for the backbone of residue 11*. *B) Both conformations of the residues shown in panel A are shown for the core mutant. Mutated residues are colored in a lighter blue. The C² of Gly69 is shown as a small sphere for clarity*. *C, D, E) Interactions between residue 6 of β-stand 1 and residue 68 of strand 5 for the wild-type ubiquitin (gray, panel C), of the core mutant (blue, panel D), and the affinity matured mutant (green, panel E). A modeled water appears linking the backbone of residue 6 with the histidine 68 side-chain in both the core mutant and WT (3ons). This interaction is directly replaced in the affinity matured mutant by a hydrogen bond between the new arginine side-chain and the backbone carbonyl of residue 6*.

The affinity matured construct has three additional mutations (R42W, Q49R, H68R). H68R sits on the final ²-strand directly adjacent to the β1β2 loop. A bridging water molecule links the carbonyl oxygen of Lys6 and the histidine side chain of residue 68 in the core mutant. This bridging water molecule is modeled in multiple previously determined WT ubiquitin crystal structures (3ons and 1yiw) and there is a corresponding unmodeled electron density peak in the original WT ubiquitin dataset (1ubq) (**Supplementary Figure 3**). In the final affinity matured construct this water-mediated interaction is replaced by a direct hydrogen bond between the introduced arginine side chain and the Lys6 carbonyl directly, stabilizing the loop in one conformation (**Figure 4 c,d,e**).

### *Conformational heterogeneity is reduced in affinity matured* ²1²3 loop *and C-terminus*

Two regions adjacent to the β1β2 loop also follow the same trend as the loop itself, displaying increased heterogeneity from the WT to the core mutant and decreased heterogeneity from the core to affinity matured mutants. The first of these areas that display heterogeneity in the core mutant is the C-terminal tail (residues 73-76), which is disordered in the electron density map of the core mutant. The density for L71R, one of the core mutations located just before the C-terminus, is not observable for the side chain beyond Cβ, but is ordered in the affinity matured structure. In the affinity matured construct, two of the three new mutations are involved in a new interaction with the C-terminus. The R42W and Q49R mutations create a cation-pi-cation stacking interaction with R72 (**Figure 5 a,b,c**). This interaction further links the C-terminus to the other β-strands of the protein, ordering residue 72 near the C-terminus.

A second area that displays significant conformational heterogeneity in the core mutant, but not in the affinity matured mutant is the loop region between the α1 helix and the β3 strand (residues 32-41). In this loop the backbone displays heterogeneity where alternative conformations are shifted by a 1.0-2.2 ², resulting from a subtle hinging at the ends of the loop (**Figure 5d**). In the most shifted region in the loop, the Asp39 backbone carbonyl, the density has two discrete peaks corresponding to two states, rather than a smooth continuum that could be modeled by a single conformation with an anisotropic B-factor. Within the α1β3 loop, we observe another pair of compensatory mutations for residues 13 (I13Y) and 34 (E34L) in the core mutant. Specifically, Tyr13 buries Leu34 (**Supplementary Figure 4 a,b**), maintaining the interaction between aliphatic groups at these sidechain positions. The hydrophilic hydroxyl group in Tyr13 is also retained in a nearly equivalent position to the carboxyl group of Glu34 in the WT protein. Although sidechain interactions appear to be maintained, this new interaction coincides with increased mobility of the α1β3 loop. Subtle differences in packing that result from these amino acid substitutions, and concomitant changes to adjacent residues, may introduce flexibility in this region. In contrast to the core mutant, the α1β3 loop in the affinity matured mutant does not display significant heterogeneity (**Figure 5e**). Although none of the affinity matured mutations are in this exact region, it is likely that the additional stabilization of the C-terminus gained from the new cation-pi-cation interaction between residues W42, R49, and R71 in the affinity matured mutant propagates to the β1β3 loop via backbone interactions. Notably, a network of hydrogen bonds connects residues 40 and 41 of the α1β3 loop with residues 70 and 72 of the C-terminus. We therefore hypothesize that the new interactions observed in the affinity matured mutant cooperate with native hydrogen bonding motifs to quench the dynamics of the α1β3 loop.

**Figure 5.**
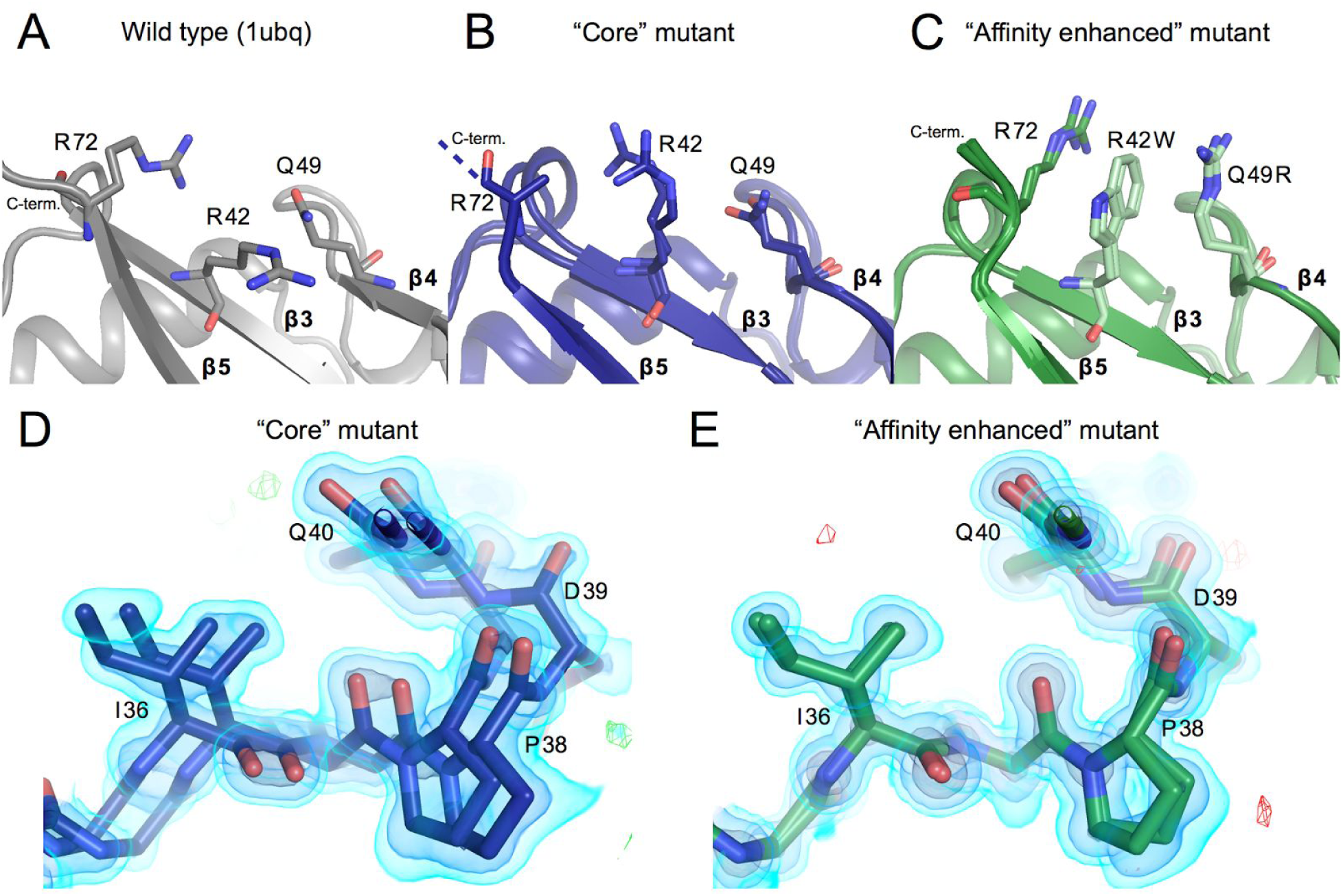
*Reduced conformational heterogeneity from core mutant to affinity matured mutant* *A, B, C) The packing of residues 42, 49, and 72 are shown. Panels and colors show the WT, core and affinity matured mutants in gray, blue and green respectively. Once mutated, residues are shown in a lighter shade of the same color. Residue Arg72 could not be fully built in the core mutant model and thus is truncated at the Cβ. Residues mutated in the affinity matured protein now show new cation-pi interactions, both between residues 72 and 42, and between 42 and 49.* *D) Residues 36-39 are shown highlighting heterogeneity in the affinity matured mutant that spans residues 32-41. There is signal for two conformations that differ in this region by a shift of as much as 2.2* *. 2FoFc map shown as a volume contoured to 3.5, 1.5 and 0.65 e ^−3^ (light blue, blue, black), with the difference FoFc map contoured to 3 e ^−3^*. *E) The heterogeneity seen in the core mutant in panel D is not seen for the affinity matured structure at the same residues. Maps are contoured as in D*.

### Both states of residue Asp52-Gly53 peptide flip occur in core and affinity matured structures

Residues Asp52 and Gly53 have been previously identified as a structural switch in ubiquitin that undergoes a discrete peptide flip (Huang et al., 2011) that exchanges on the microsecond timescale (Sidhu et al., 2011; Smith et al., 2016). The original crystal structure of wild-type ubiquitin (1ubq), a standard in computational benchmarking studies, shows the peptide in the “NH-out” state with the Asp52 carbonyl making a hydrogen bond to the backbone of the β1-helix starting residue Glu24. In the crystal structure (3ons) from conditions similar used for solid state NMR the “NH-in” state is seen, where the Glu24 side chain is swung down relative to the NH-out conformation to make a hydrogen bond with the Glu24 backbone and the G53 NH group. This observation has provided a structural rationale for slow NMR dynamics measurements in solution (Majumdar and Ghose, 2004; Massi et al., 2005) and in solid state (Tollinger et al., 2012): the chemical shifts of backbone amides surrounding residue 24 are perturbed as the sidechain transitions to a new rotamer. Moreover, this flip is thought to be a key structural switch between different states of ubiquitin. The flip state of the peptide can be predicted from the backbone coordinates of other residues clear across the protein (Smith et al., 2016).

The major conformation in both of our structures corresponds to the NH-in conformation found in the 3ons structure. When modeling this region as a single conformer, we observed signals in the Fo-Fc map supporting an alternative conformation corresponding to the NH-out state (**Figure 6a**). In our final model, the peptide flip is modeled in both states (**Figure 6b**) for both structures at occupancies of 60-70% for the major NH-in and 30-40% for the NH-out.

**Figure 6.**
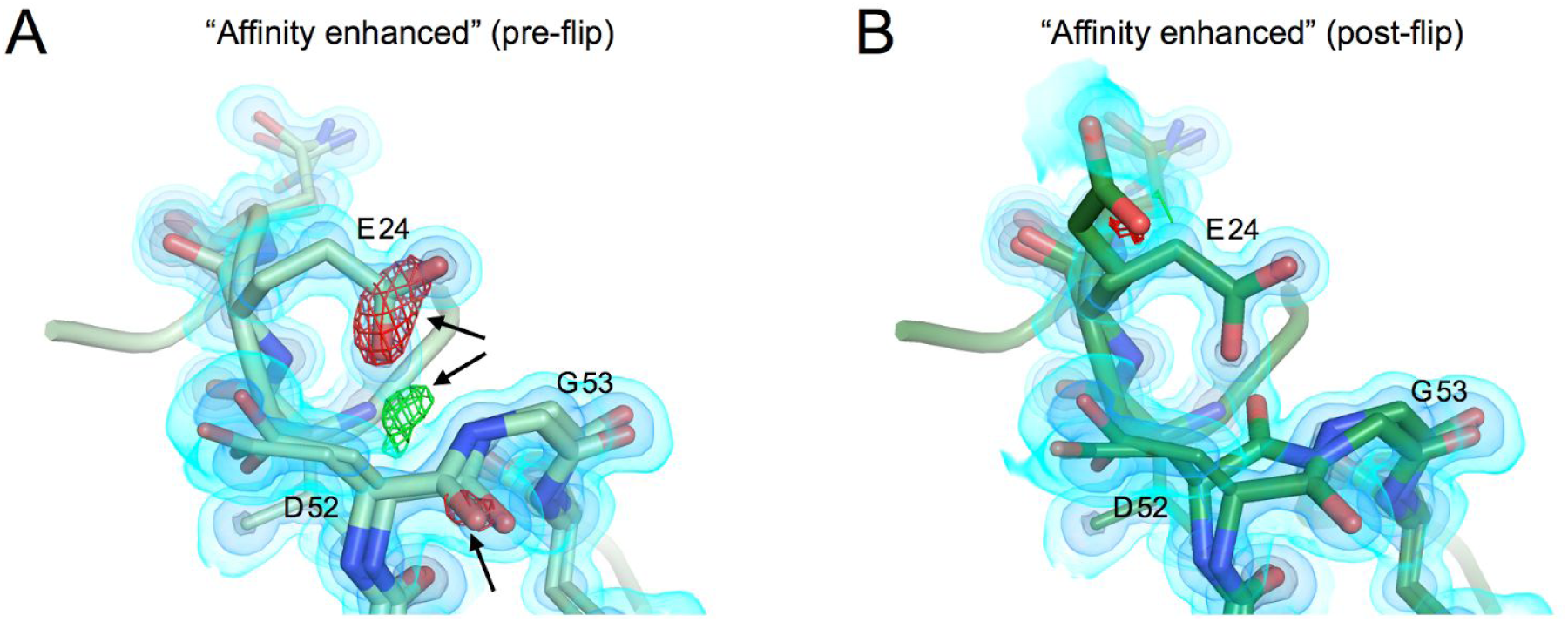
*Asp52-Gly53 peptide flip is seen in both states* *A) Model and maps from the affinity matured mutation prior to modeling a peptide flip in this region. 2FoFc map shown as a volume contoured to 3.5, 1.5 and 0.65 e ^−3^ (light blue, blue, black), with the difference FoFc map contoured to 3 e ^−3^. There are clear difference features both positive (green) and negative (red) highlighted by black arrows*. *B) Modeled peptide flip in final structure of the core mutant. Maps are contoured as in A. The difference features are now gone*.

Interestingly, the NH-in state has been implicated in the binding mode of ubiquitin to the USP class of deubiquitinases which includes the target, USP7 (Smith et al., 2016). While the population of the NH-in state may have been increased relative to WT, it remains surprising that both peptide flip states are observed in our structures given the strong association of this peptide flip with the binding of USP7. These structures indicate that it is possible for both of these states to exist within the same crystal form, and provide an additional example of how multiconformer X-ray models can be used to provide a structural basis for dynamics observed by NMR.

## Discussion

We have structurally characterized two ubiquitin variants created via a combination of computational protein design calculations and phage display. These variants are poorly modeled by traditional single conformer models, or even with existing automated model building tools for regions of high heterogeneity. Enabled by room temperature X-ray data collection, and manually-built multiconfomer models, we have described the emergence and quenching of conformational heterogeneity along a protein design and engineering trajectory. From these multiconformer models, we observed the interplay of computational protein design and laboratory evolution, describing both the core and affinity matured constructs that were developed to bind tightly to USP7. Notably, the intermediate core mutant displayed significant conformational heterogeneity across the majority of the protein, varying in magnitude. The large-scale backbone motions of the β1β2 loop are in direct contrast to the original goal of stabilizing a single pro-USP7 binding conformation. Instead, the core mutations enhance the flexibility of the protein, creating β1β2 loop conformations that are distinct from states in the hinge-like motion predicted by NMR analyses of the WT protein (Lange et al., 2008). In contrast, additional surface mutations introduced in the final affinity matured construct cooperate to reduce flexibility and the β1β2 loop can be modeled in a single conformation.

Our observations suggest that the process of rational protein design followed by directed evolution resembles simulated annealing procedures used to escape local conformational minima in X-ray refinement (Brünger et al., 1997; Korostelev et al., 2002). For these ubiquitin variants, the core mutations, selected based on Rosetta calculations and phage display, act like the heating step. These mutations disrupt the natural dynamics and packing of the WT protein, creating a large and diverse ensemble of states. Relative to the initial WT conformation, these dynamics likely increase sampling of states that have the desired function; however, many undesirable conformations are also sampled. Further affinity maturation acts like the cooling step in simulated annealing, selectively stabilizing the functional states. This final pattern of flexibility changes is similar to what has been hypothesized for designed enzymes subjected to directed evolution (Bhabha et al., 2015; Kiss et al., 2013) and observed in antibody maturation (Adhikary et al., 2015; Jimenez et al., 2004). The discordant mapping between the conformational landscape, which is most heterogeneous in the “core” mutant, and functional landscape, which shows the largest functional gain from the initial “core” phage display, demonstrates the complex interplay between conformational dynamics and function. Although our observations are in the context of the evolution of new binding specificity, they can likely be translated to the more complicated challenge of evolving new catalytic activities (Campbell et al., 2016)

The ability for proteins to evolve new functions in this way relies on the fact that the hydrophobic cores of most proteins can accommodate many alternative sequences without compromising stability (Lim and Sauer, 1989). This permissiveness can be exploited to alter functional specificity (Koulechova et al., 2015). Here, we observed how changes in side chain packing between mutations in direct contact can lead to dramatic changes in backbone flexibility. Therefore, core mutations, even with multiple compensatory mutations, can disrupt the natural dynamic packing and may increase backbone flexibility. These changes lead to altered dynamics, which can be exploited for evolving new functions. The nearly global quenching of backbone heterogeneity in the affinity matured mutant, both in regions directly adjacent to mutations and across the protein, point to the importance of the cooperativity in these dynamic packing interactions.

Surprisingly, the reduced dynamics in the affinity matured variant are enabled by introduction of new surface interactions, including a cation-pi-cation interaction observed between residues 72, 42, and 49. These new side chains apparently cooperate to “freeze” motion on the backbone behind these new interactions. The importance of this cation-pi-cation motif in this structure suggests possibilities to improve rational design of new or altered protein function by incorporating rarer interactions in the design process. Despite efforts to develop potentials for cation-pi interactions in Rosetta (Misura et al., 2004), they are not currently modeled in the standard Rosetta energy function and thus would be completely missed in the design process. Cation-pi interactions have recently been used to stabilize a miniature designed protein (Craven et al., 2016) and are common in naturally occurring proteins (Dougherty, 2013). In addition, ordered waters, which are not directly accounted for by Rosetta (Jiang et al., 2005), play a key role in the stabilization of hydrogen bond networks such as between the WT His68 and the β1β2 loop. These overlooked features of protein structure play key roles in conformational stabilization and could be incorporated to improve the protein design process.

Interestingly, despite the dramatic increase in affinity for the binding partner USP7, the β1β2 loop in our apo-crystal structures does not adopt the “down” conformation seen in the WT ubiquitin USP7 complex. Instead, the affinity matured loop state resembles the WT apo “up” state (**Figure 3d**). It is unclear whether the “down” conformation is stabilized by these mutations, albeit still as a minor conformation that cannot be detected crystallographically. Rather than disrupting the dynamics of the β1β2 loop in such a way that the “down” state is the only state accessed, one explanation for the enhanced affinity to USP7 may be that the mutations introduced key interfacial residues that produce a more complementary surface to the large binding cleft of USP7. Alternatively, the mutations may reduce the energetic penalty for reorganizing into the bound conformation in an induced fit mechanism. Future studies of the core and affinity matured mutants bound to USP7 are needed to answer these questions. Given the recent interest in using ubiquitin variants for structural biology chaperones as *in vitro* modulators of the ubiquitin proteasome system (Canny et al., 2016; Ernst et al., 2013; Gorelik et al., 2016; Phillips et al., 2013; Zhang et al., 2016), there may be additional opportunities to use these proteins to learn about the importance of conformational dynamics in protein function (Phillips et al., 2013).

## Experimental Procedures

### Protein purification

Protocol adapted from Zhang et al ((Canny et al., 2016; Ernst et al., 2013; Gorelik et al., 2016; Phillips et al., 2013; Zhang et al., 2016)) and references therein. Each of the ubiquitin constructs were expressed from a pET derivative vector containing the protein gene with a TEV cleavable N-terminal His6 tag in *E. coli* BL21(DE3) cells. Cells were grown in LB media to an OD600 of 0.6-0.8, and were then induced with 1mM IPTG at 37 °C for 4 hours. Cell pellets were resuspended into 25 mM Tris pH 8.0, 300 mM NaCl and protease inhibitors. The resuspended pellet was then lysed by Emusiflex (Avestin). Lysate was cleared by centrifugation at 15,000 rpm. Supernatant was flowed over a 5 mL Ni-NTA column. Ni-NTA column was washed with 20 mM Tris pH 7.0, 300 mM NaCl, 20 mM imidazole and eluted with 20 mM Tris pH 7.0, 300mM NaCl, 500 mM imidazole. Elution was dialyzed into 20 mM Tris pH 8.0, 150 mM NaCl and then cleaved with TEV overnight at 4 °C. The sample was then loaded back onto a Ni-NTA column, and the flow through was collected. Sample was then concentrated to less than 10 mL and loaded onto a S75 column equilibrated with the previous buffer (20 mM Tris pH 8.0, 150 mM NaCl). Protein concentration of the pooled fractions was assessed after concentration down to 1-2 ml via a BSA assay and monitored during further concentration via absorption. Protein was stored only overnight at 4 °C before concentration and use for crystallization.

### Crystallization and data collection

Protein solution from was concentrated to 20 mg/ml for the core mutant and 10 mg/ml for the affinity matured mutant. Then 1ul of protein solution was mixed with 1ul of precipitation solution in hanging drop trays. Precipitation solution for core mutant contained 0.1 M citric acid pH 4.6, and 2.6 M ammonium sulfate. For the affinity matured mutant, the precipitation solution contained 0.1 M citric acid pH 4.2, and 2.2 M ammonium sulfate. Crystals formed overnight and grew to full size over the course of several days.

### Optimizing the diffraction resolution cutoff for multiconformer modeling

After optimizing crystals for the room temperature collection of high resolution datasets for the initial core and the final affinity matured constructs, we had to select an appropriate resolution cutoff for our data. Because the ability to detect and faithfully model conformational heterogeneity is resolution dependent, we aimed to push the resolution of our dataset as far as possible. High resolution reflections, despite their lower signal-to-noise, can contain meaningful information that can improve the map and model, even past more traditional methods of picking a resolution cutoff. Instead of simply using CC1/2, completeness and I/β, we used the Karplus and Diederichs approach of monitoring Rfree in parallel refinements to determine the optimal resolution cutoff for our datasets (Karplus and Diederichs, 2012). When using parallel refinements to determine the optimal resolution cutoff, additional higher resolution reflections are judged to contain meaningful signals only if model agreement increases in lower resolution bins after refinement.

We initially chose a conservative resolution cutoff of 1.16 Å to begin our parallel refinement tests, and created additional bins of reflections to be added in 0.04 Å increments up to a high resolution cutoff of 0.96 Å. While a 0.04 Å resolution change may seem small, ~2000 unique reflections are added in each bin. The values for CC1/2 in the high resolution bins remained over 50% up to 1.04 Å resolution, while completeness began to drop at resolutions better than 1.12 Å (**Table 1**). To begin the resolution test, first we built a single conformer model using the reflections up to the first cutoff (1.16 Å). We then built alternative conformations into strong difference density signals corresponding to clear alternative conformations. At this point, we added reflections from each additional resolution bin ranging from 1.16 Å to 0.96 Å, and re-refined the model to convergence in parallel refinements. Based on the R-values from these parallel refinements, the determined optimal resolution cutoffs were 1.12 Å for the core mutant and 1.08 Å for the affinity matured construct (**Table 1**, **Supplemental Figure 6**).

**TABLE 1:** *Data statistics at different resolution cutoffs* A) Table of values for core mutant structures at different maximal resolutions. Completeness, CC1/2 and I/σ are shown for the high resolution bin at each resolution cutoff choice. Both Rwork and Rfree percent values up to resolution cutoff shown for each row corresponding to the structure built using data to resolution cutoff shown for each column. Each column represents different structures built using data with different resolution cutoffs, however, structures can only be compared when the same reflections are compared, i.e. values in the same row. B) Statistics for affinity matured structures at different resolution cutoffs. As above, completeness, CC1/2 and I/σ are shown for the highest resolution bin, and R/Rfree statistics are shown for all reflections within the designated resolution cutoff (Rows).

| A - Core Mutant |  |  |  |  |  |  |  |  |
| --- | --- | --- | --- | --- | --- | --- | --- | --- |
| Completeness (%) | CC1/2 (%) | Rmerge (%) | I/sigma | Refinement Resolution<br>R-factor calculation Resolution | 1.16 Å | 1.12 Å | 1.08 Å | 1.04 Å |
| 100 | 96.8 | 17.9 | 8.46 | 1.16 Å | 14.4/16.3 | **14.2 / 16.0** | 15.1 / 16.2 | 15.0 / 16.1 |
| 100 | 95.3 | 22.4 | 6.89 | 1.12 Å |  | **14.2 / 16.1** | 15.1 / 16.3 | 15.0 / 16.4 |
| 97.2 | 89.4 | 29.9 | 4.47 | 1.08 Å |  |  | 15.1 / 16.5 | 15.1 / 16.5 |
| 91.1 | 74.7 | 41.8 | 2.41 | 1.04 Å |  |  |  | 15.2 / 16.7 |

**TABLE 1: Data statistics at different resolution cutoffs**
| B - Affinity Matured Mutant |  |  |  |  |  |  |  |  |  |  |
| --- | --- | --- | --- | --- | --- | --- | --- | --- | --- | --- |
| Completeness (%) | CC1/2 (%) | Rmerge (%) | I/sigma | Refinement Resolution<br>R-factor calculation Resolution | 1.16 Å | 1.12 Å | 1.08 Å | 1.04 Å | 1.00 Å | 0.96 Å |
| 100 | 98.1 | 24.8 | 9.24 | 1.16 Å | 11.0 / 12.5 | 11.1 / 12.4 | **11.1 / 12.3** | 11.3 / 12.4 | 11.4 / 12.4 | 11.6 / 12.6 |
| 100 | 96.9 | 32.5 | 7.01 | 1.12 Å |  | 11.0 / 12.5 | **11.1 / 12.4** | 11.2 / 12.5 | 11.3 / 12.5 | 11.5 / 12.6 |
| 96.8 | 90.9 | 47.9 | 4.22 | 1.08 Å |  |  | **11.2 / 12.5** | 11.3 / 12.5 | 11.3 / 12.6 | 11.5 / 12.7 |
| 87.8 | 76.7 | 66.6 | 2.28 | 1.04 Å |  |  |  | 11.5 / 12.8 | 11.5 / 12.8 | 11.7 / 12.9 |
| 68.7 | 41.3 | 118.3 | 1.00 | 1.00 Å |  |  |  |  | 11.8 / 13.1 | 12.0 / 13.3 |
| 34.3 | 7.7 | 233.1 | 0.38 | 0.96 Å |  |  |  |  |  | 12.2 / 13.5 |

## Author Contributions

Performed experiments JTB

Performed refinements JTB, MCT

Designed experiments JTB, CNC, JEC, JSF

Wrote manuscript JTB, JSF

Edited manuscript JTB, MCT, CNC, JEC, JSF

## Acknowledgements

JTB is supported by a NSF GRFP. M.C.T. is supported by NIH F32 HL129989. J.S.F. is supported by a Searle Scholar Award from the Kinship Foundation, a Pew Scholar Award from the Pew Charitable Trusts, a Packard Fellowship from the David and Lucile Packard Foundation, NIH DP5 OD009180, NIH R21 GM110580, and NSF STC-1231306. Data collection at BL831 at the Advanced Light Sources is supported by the Director, Office of Science, Office of Basic Energy Sciences, of the U.S. Department of Energy under Contract No. DE-AC02-05CH11231, UC Office of the President, Multicampus Research Programs and Initiatives grant MR-15-32859, and the Program Breakthrough Biomedical Research, which is partially funded by the Sandler Foundation. We acknowledge helpful conversations with H. van den Bedem, T. Kortemme, N. Betheil and the Fraser lab, especially D. Keedy.

## Accession Numbers

5tof - Core mutant (U7Ub25)

5tog - Affinity matured mutant (U7Ub25.2540)

## Supplementary Figures

**Sup. Table 1.** *Data collection and refinement statistics*.

|  | **Core (u7ub25)** | **Affinity matured (u7ub25_2540)** |
| --- | --- | --- |
| **Wavelength** | 0.9537 | 0.9537 |
| **Resolution range** | 22.04 - 1.12 (1.16 - 1.12) | 38.04 - 1.08 (1.119 - 1.08) |
| **Space group** | P 31 2 1 | P 31 2 1 |
| **Unit cell** | 42.72 42.72 54.88 90 90 120 | 43.92 43.92 55.44 90 90 120 |
| **Total reflections** | 130992 (13152) | 311084 (26130) |
| **Unique reflections** | 22427 (2242) | 27011 (2620) |
| **Multiplicity** | 5.8 (5.9) | 11.5 (9.9) |
| **Completeness (%)** | 0.98 (1.00) | 1.00 (0.98) |
| **Mean I/sigma(I)** | 18.25 (7.13) | 28.21 (4.70) |
| **Wilson B-factor** | 9.91 | 8.95 |
| **R-merge** | 0.05401 (0.2237) | 0.04694 (0.454) |
| **R-meas** | 0.05949 (0.2458) | 0.0492 (0.4788) |
| **CC1/2** | 0.998 (0.955) | 1 (0.929) |
| **CC\*** | 1 (0.988) | 1 (0.982) |
| **Reflections used in refinement** | 22426 (2242) | 27011 (2622) |
| **Reflections used for R-free** | 1371 (137) | 1574 (155) |
| **R-work** | 0.1543 (0.1480) | 0.1032 (0.1184) |
| **R-free** | 0.1752 (0.2043) | 0.1212 (0.1568) |
| **CC(work)** | 0.943 (0.946) | 0.976 (0.973) |
| **CC(free)** | 0.914 (0.892) | 0.981 (0.953) |
| **Number of non-hydrogen atoms** | 1135 | 1358 |
| **macromolecules** | 1093 | 1247 |
| **ligands** | 5 | 10 |
| **solvent** | 37 | 101 |
| **Protein residues** | 73 | 75 |
| **RMS(bonds)** | 0.016 | 0.009 |
| **RMS(angles)** | 1.57 | 1.05 |
| **Ramachandran favored (%)** | 99 | 100 |
| **Ramachandran allowed (%)** | 1.4 | 0 |
| **Ramachandran outliers (%)** | 0 | 0 |
| **Rotamer outliers (%)** | 1.71 | 0 |
| **Clashscore** | 2.71 | 1.18 |
| **Average B-factor** | 12.62 | 10.95 |
| **macromolecules** | 12.27 | 10.02 |
| **ligands** | 21.76 | 15.89 |
| **solvent** | 21.56 | 21.92 |

Statistics for the highest-resolution shell are shown in parentheses.

**Sup. Figure 1:**
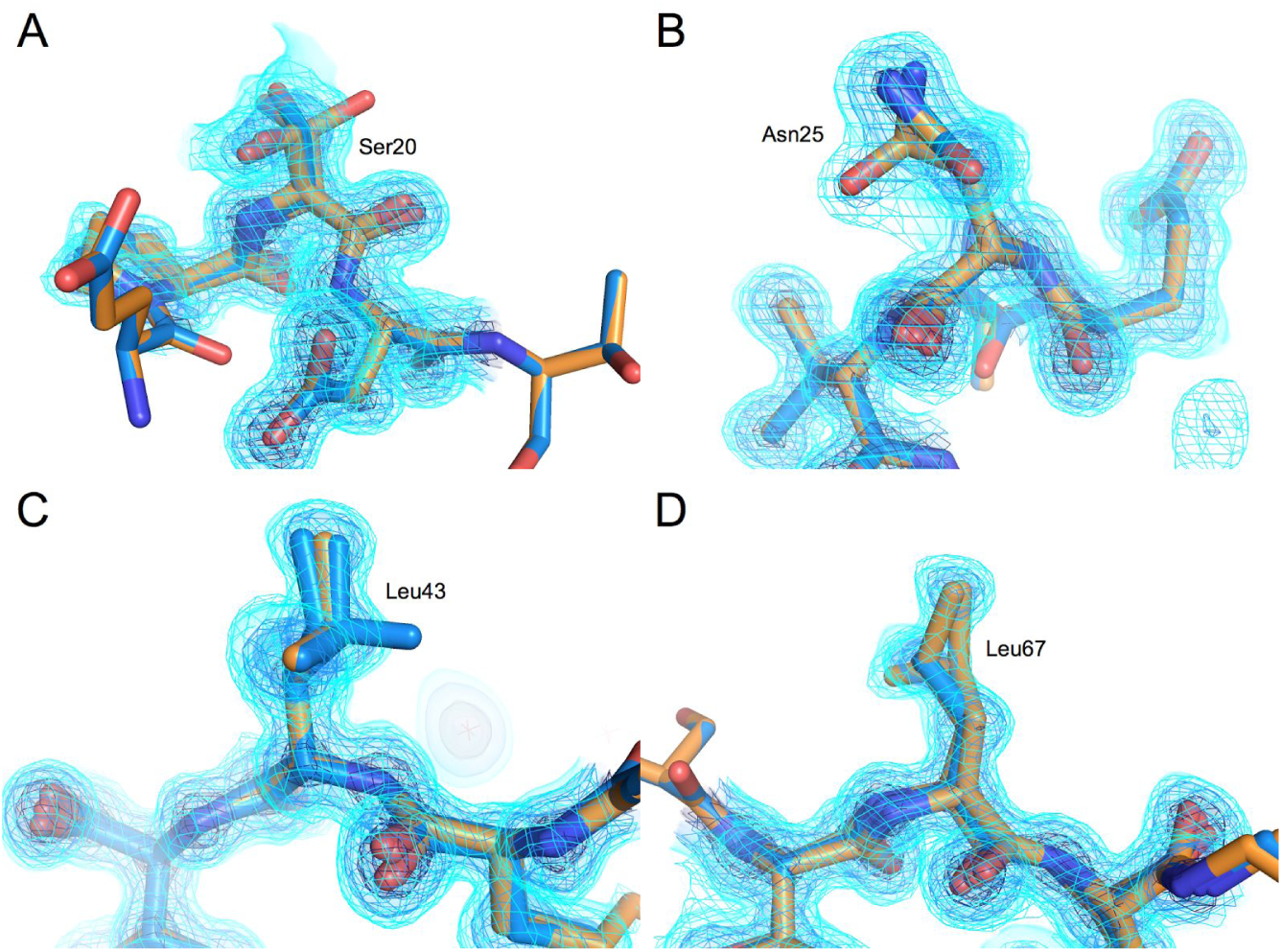
*Regions of interpretative difference between MCT and JTB blinded-models*. *Models built by JTB (blue) and MCT (orange) are both shown overlayed. The 2Fo-Fc density map is contoured to 0.65 eÅ^−3^, 1.5 eÅ^−3^ and 3.5 eÅ^−3^ (lightest to darkest); JTB maps are shown as volume, and MCT maps shown as mesh*. *A) Residue Ser20 shown, where MCT has build a third conformer of the serine side-chain that is not strongly supported by the density, and was deleted in the final model (not shown here)*. *B) Residue Arg25 where MCT built one extra conformation of asparagine side-chain supported by the density. This conformer was incorporated into final model*. *C) Here, model built by JTB has an extra Leu43 conformer that sticks out of the density*. *D) An extra Leu67 side-chain was built by MCT that was later interpreted as a different two-conformer model into signal that became more clear after further refinement*.

**Sup. Figure 2:**
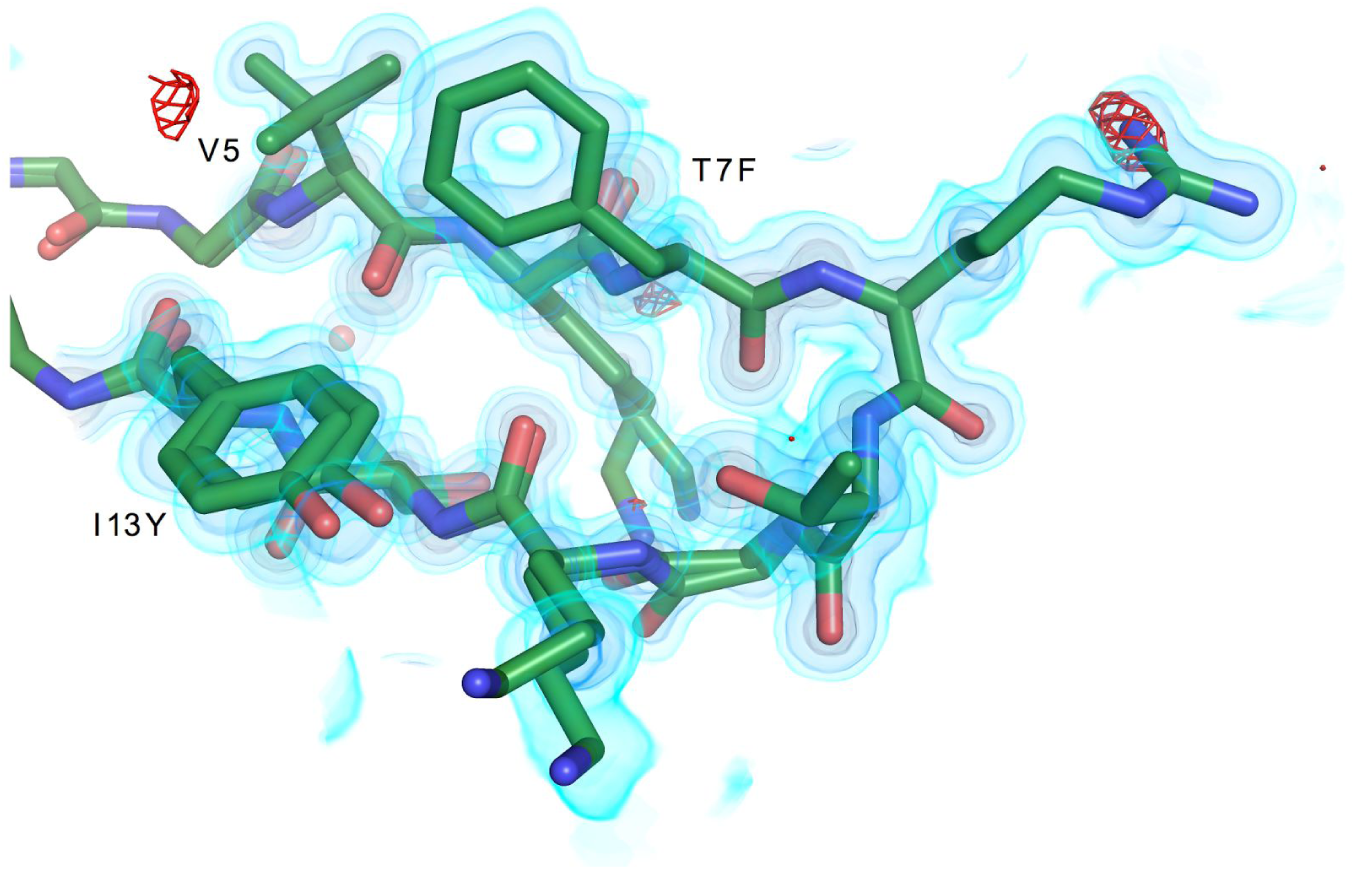
*Affinity matured β1β2 loop with density*. *Residues 5-13 shown as sticks, with 2Fo-Fc density map contoured to 0.65 eÅ^−3^, 1.5 eÅ^−3^ and 3.5 eÅ^−3^ from lightest to darkest. Fo-Fc difference map contoured at 3 e ^−3^*

**Sup. Figure 3:**
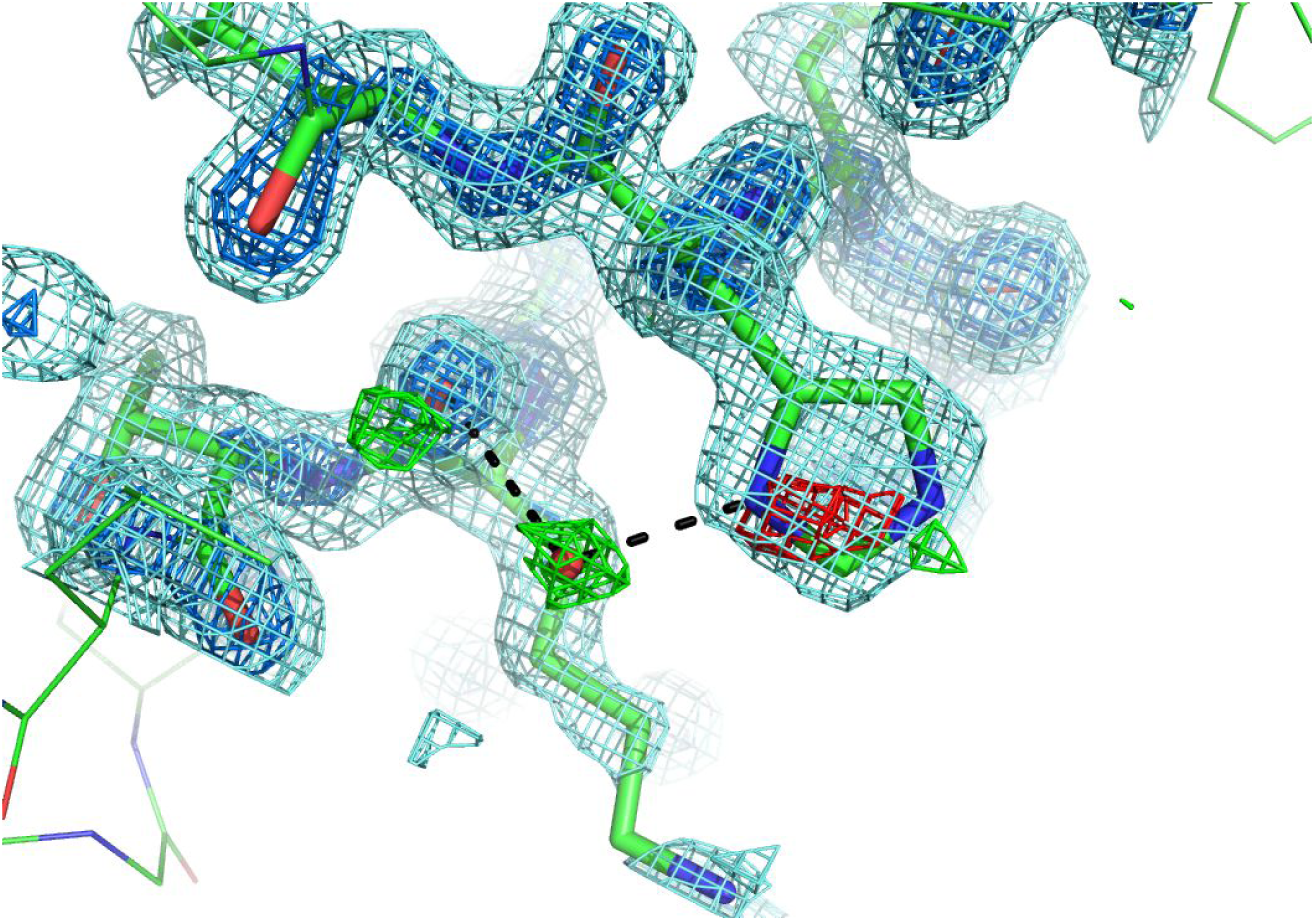
*Unmodeled water bridging His68 and Lys6*.

**Sup. Figure 4:**
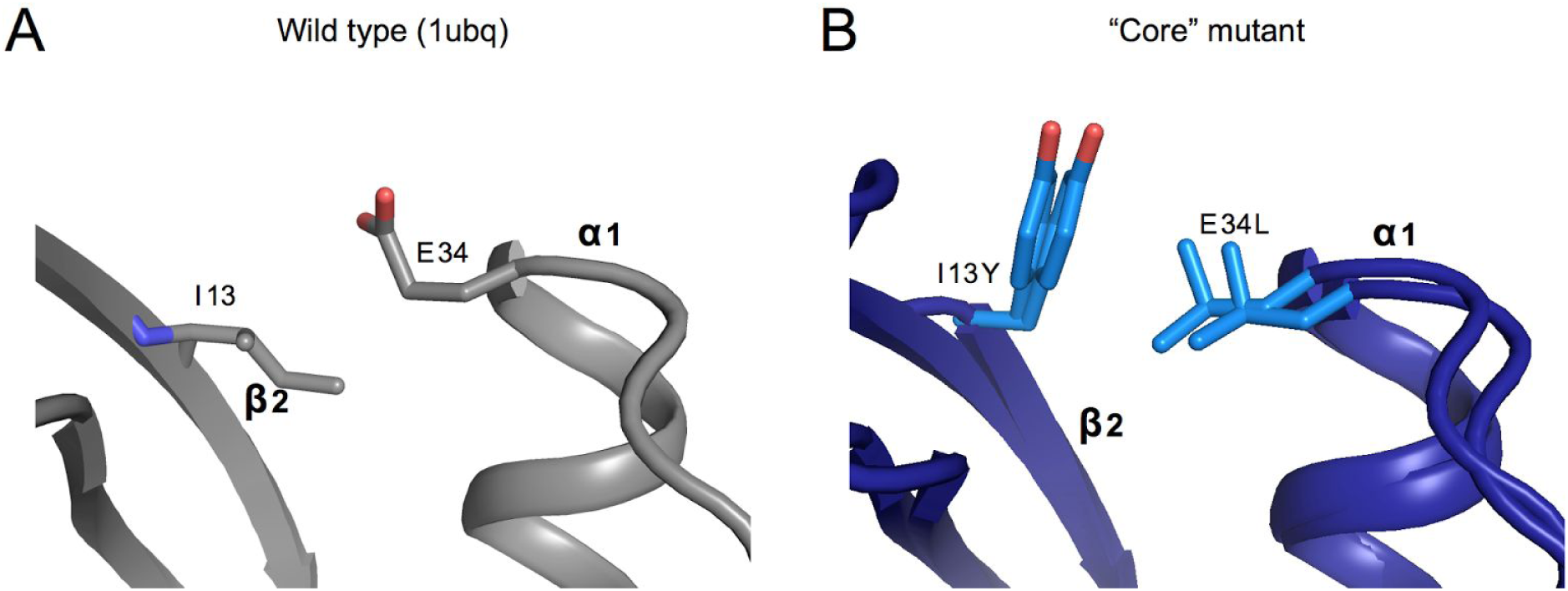
*Compensatory mutation at residues E34L and I13Y* *D) Packing interaction between the* β*2-strand’s I13 with the helix cap residue Glu34*. *E) Altered packing interaction of residues 13 and 34 in the core mutant. Both conformations of the residues are shown, and the mutant side-chains are shown in a lighter blue*. *WT ubiquitin from crystal structure 1ubq, showing positive difference density for a water (built here) bridging the His68 side-chain and the Lys6 carbonyl. Hydrogen bonds linking atoms to built water are shown as dashed lines. Residues 5-7 and 67-69 are shown as sticks. The 2Fo-Fc map is contoured at 0.65 eÅ^−3^, 1.2 eÅ^−3^, from light to dark blue. The Fo-Fc map is contoured at 0.25 eÅ^−3^*.

**Sup Figure 5:**
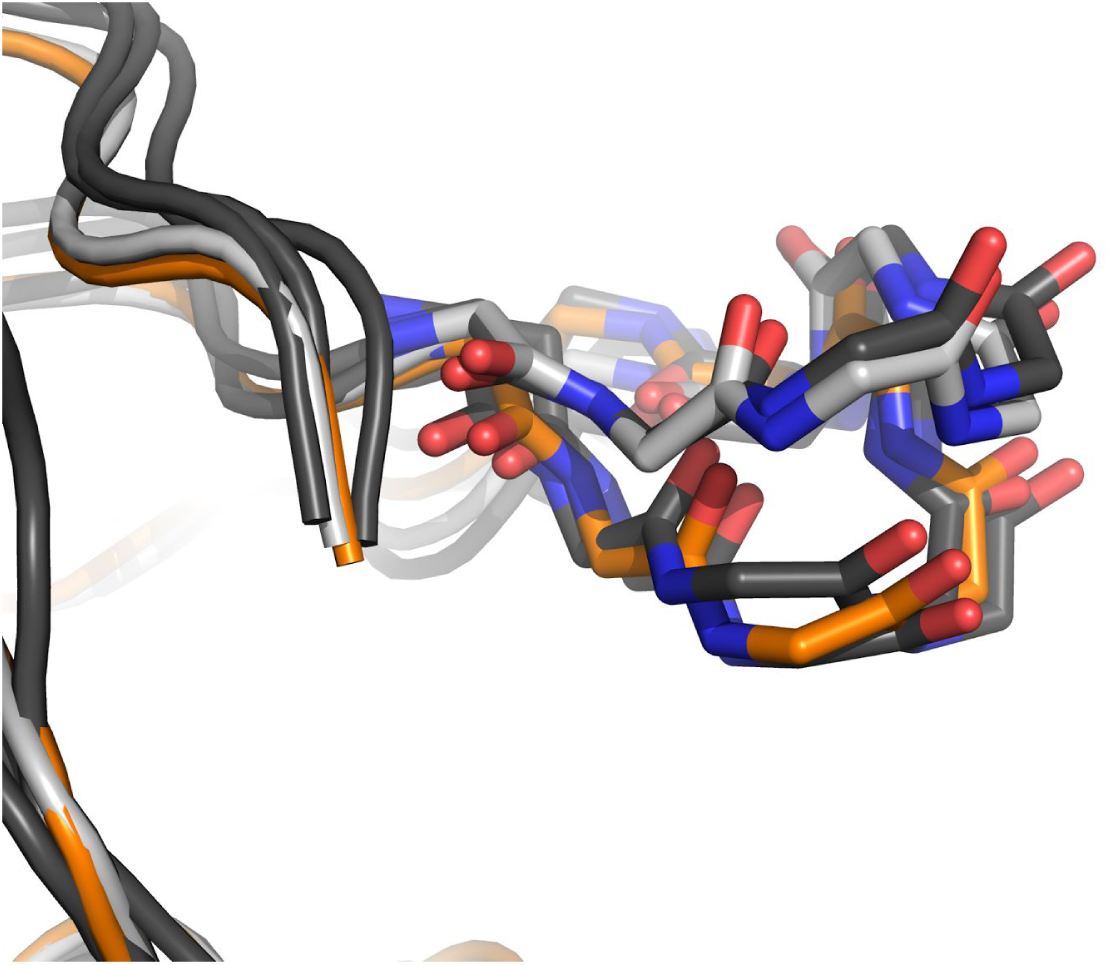
*Overlay of ubiquitin models highlighting the β1β2 loop’s up and down conformations. The WT model from 3ons (light gray) is overlaid with the USP7 bound WT ubiquitin from 1nbf (orange) with most similar states in the 2k39 ensemble (dark gray)*.

**Sup. Figure 6:**
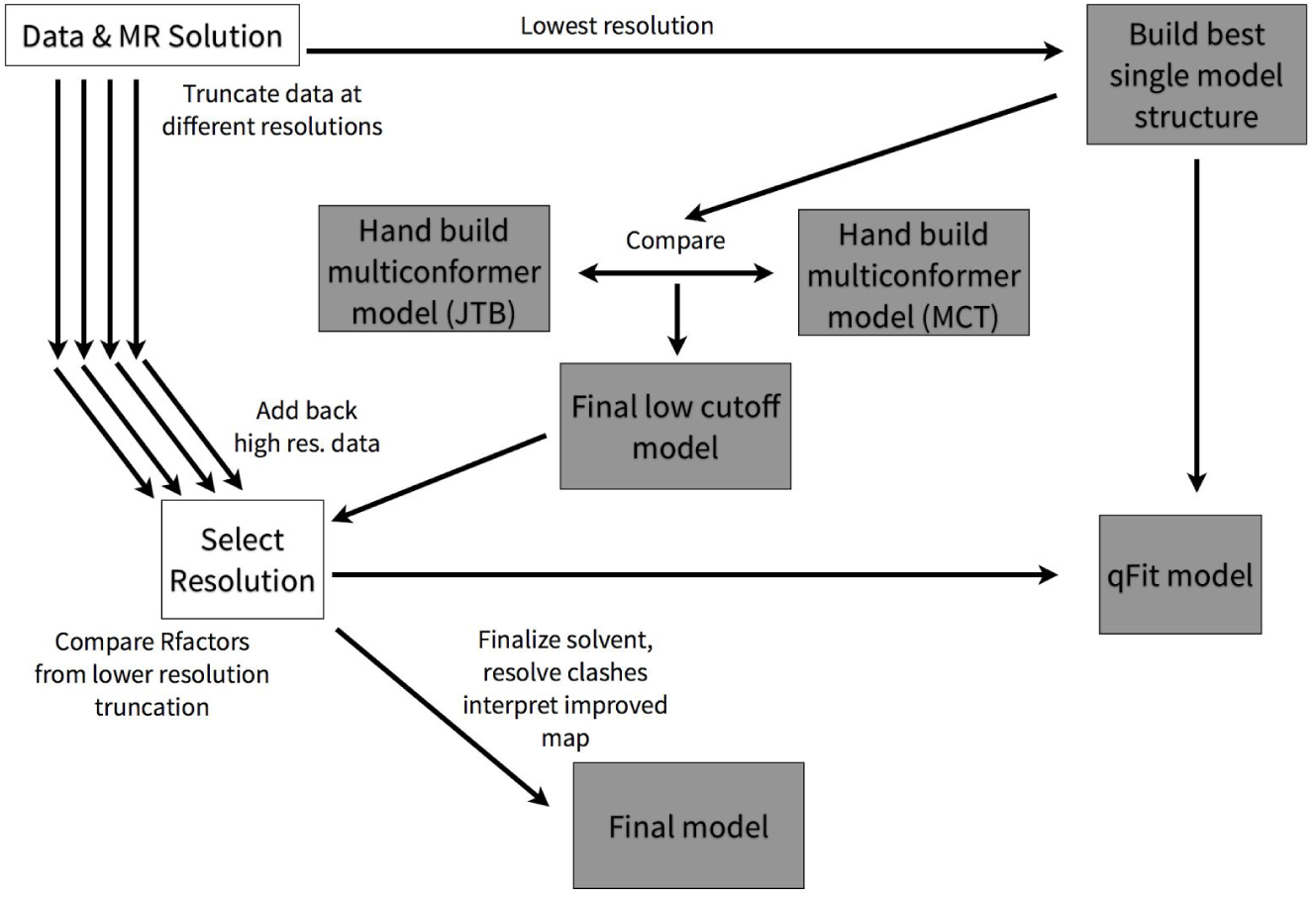
*Chart illustrating strategy for selecting a resolution cutoff and refining a final model. After processing raw images using the diffraction data processing program XDS (Kabsch, 2010) at the highest resolution (0.96), data was truncated to several lower resolution thresholds. Reflections selected for the Rfree set were created for the highest resolution data are consistent across all resolution cutoffs. The lowest resolution truncation was fed into molecular replacement with WT ubiquitin (PDB ID: 1ubq) used as a search model. After a single conformer model was built, the additional alternative conformations were identified by both JTB and MCT authors independently. After comparing JTB and MCT models, a final model (at 1.16) was fed into parallel refinements against the higher resolution data for each resolution cutoff. Rwork and Rfree were compared to select best resolution for which to move forward. This final resolution was used to feed the single conformer model into automated model building by qFit. The final model determined using the reflections up to 1.16 was further interpreted by making additional solvent changes, resolving clashes and inspecting signal resolved by the higher resolution data*.

